# Associations between maternal pre-pregnancy BMI and infant striatal mean diffusivity

**DOI:** 10.1101/2023.09.11.557121

**Authors:** Aylin Rosberg, Harri Merisaari, John D. Lewis, Niloofar Hashempour, Minna Lukkarinen, Jerod M. Rasmussen, Noora M. Scheinin, Linnea Karlsson, Hasse Karlsson, Jetro J. Tuulari

## Abstract

**Background/Objectives:** It is well-established that parental obesity is a strong risk factor for associates with offspring obesity. Further, a converging body of evidence now suggests that maternal weight profiles may affect the developing offspring brain in a manner that confers future obesity risk. Here, we investigated how pre-pregnancy maternal weight status influences the reward-related striatal areas of the offspring brain during *in utero* development.

**Methods:** We used diffusion tensor imaging to quantify the microstructure of the striatal brain regions of interest in neonates (N = 116 mean gestational weeks at birth 39.88, SD = 1.14; and at scan 43.56, SD = 1.05). Linear regression was used to test the associations between maternal pre-pregnancy body mass index and infant striatal mean diffusivity.

**Results:** A strong positive association was found between the maternal pre-pregnancy body mass index and newborn left caudate nucleus mean diffusivity. Results remained unchanged after the adjustment for covariates.

**Conclusions:** *In utero* exposure to maternal adiposity might have a growth impairing impact on the mean diffusivity of infant left caudate nucleus. Considering the involvement of caudate nucleus in regulating eating behaviour and food-related reward processing later in life, this finding calls for further investigations to define the prognostic relevance of early life caudate development and weight trajectories of the offspring.

## 1. Introduction

Parental obesity plays a critical role in the mental and physical health of the offspring in many stages of their lives. The existing body of research indicates that if at least one parent is obese, regardless of the children’s weight status or age interval, the children are at high risk of obesity as early as at age two^1^. The risk is higher if the mother is obese compared to the father being obese ^2^.

One of the most accepted explanations for the influence of high maternal adiposity on the offspring weight status has been the genetic heritability. The implicated genes are mainly involved in the neural networks controlling eating habits such as the caloric intake, appetite regulation and even compulsive hyperphagia ^3,4^. However, the predictive power of genetics fluctuates over the lifespan of the offspring ^5^ and it does not account for the greater relation between obesity in the offspring and maternal obesity compared to paternal obesity. In addition, the environmental factors or other behavioural traits are also known to affect adult body mass index (BMI) ^6^. This supports the assertion that obesity is a “heritable neurobehavioural disorder that is highly sensitive to environmental conditions”, including *in utero* exposures.

Striatum (caudate nucleus and lentiform nucleus that comprises putamen and globus pallidus) has recurrently been found to be involved in hedonic eating, food related motivation and reward processes ^7–10^. Differences have been found in neural processing of unhealthy, *i.e.* high-sugar content, food between healthy weight and overweight or obese individuals ^11,12^. Obesity is associated with structural changes in striatum in the adolescent ^13^ and adult population ^14–16^. Moreover, the functional and structural changes in the striatum may predict future weight gain^17–20^.

Taken the paramount role of maternal prenatal factors that affect *in utero* brain development, it is crucial to investigate the neurodevelopment of the offspring who were prenatally exposed to maternal overweight/obesity. Still, there is only limited research investigating the infant neurodevelopment of those who were exposed to prenatal maternal obesity. Two extant studies have shown that the altered functional connectivity in reward processing and cognitive control, similarly to that observed in obese adults, was also observed in 2-week-old infants with mothers who had high BMI ^21,22^. In addition, decreased white matter integrity ^23^ and increased mean diffusivity in hypothalamus ^24^ were observed in infants born to mothers with high BMI. Diffusion tensor imaging (DTI) has been an indispensable method to study the white matter, since it provides quantitative scalar values to assess the microstructural integrity. Thanks to the recent technological advancements, it has become possible to accurately study the microstructure of the gray matter as well ^25^, including the microstructural integrity of the subcortical regions ^26–28^. One of the four scalars derived from DTI images is mean diffusivity (MD). MD is the measure of free molecular motion of water in the tissue, therefore, regarded as the most suitable scalar for assessing gray matter structures ^29^. It represents the tissue density and reflects the overall ratio water volume between intra/extracellular tissue compartments ^30^. The free-water fraction in the tissue is positively associated with the MD value and negatively associated with presence of more cellular structures. Therefore, decreased MD has been interpreted as a marker of high tissue density ^31^. In neonates, MD is one of the most used DTI scalars and myelination is suggested as a likely cause for the observed decrease in MD after birth ^32^.

In this study, we investigated the association between the maternal pre-pregnancy BMI of the mothers and the neurodevelopmental maturation of striatum in their infants as expressed by MD. We hypothesized that there would be a positive relationship between the maternal pre-pregnancy BMI and MD values of the striatum in the newborn infant brain.

## 2. Subjects and Methods

### 2.1. Study Data

The data were collected as a part of the FinnBrain Birth Cohort Study whose broad goal is to study the effects of genes and environment on the developmental and mental health of children^33^.

The study included the data of 180 infants who were scanned between from 2 to 5 weeks of age (counted from the estimated due date), *i.e.,* 1 – 6 weeks after birth in years between 2012 – 2016. Inclusion criterion of the scanning sessions for infants was being born after gestational week 35 from singleton pregnancies. The exclusion criteria were occurrence of any perinatal complications with potential neurological consequences and being previously diagnosed with a central nervous system anomaly or an abnormal finding in a previous magnetic resonance imaging (MRI) scan with clinical indications ^34^. After the neuroimaging data preprocessing and related quality control, neuroimaging data of 122 infants that had successful structural and diffusion MRI data were available for the further analyses ^35^.

Obstetric data and maternal demographics were retrieved from the Finnish Medical Birth Register of the National Institute for Health and Welfare (http://www.thl.fi). The mothers had no history of alcohol or drug abuse, severe psychiatric disorders, epilepsy or related medication use during pregnancy. Overall, the data of 116 mother-infant dyads were used in the statistical analyses.

### 2.2. Image Acquisition

Imaging was performed during natural sleep ^35^. MRI data were acquired using a Siemens Magnetom Verio 3T scanner (Siemens Medical Solutions, Erlangen, Germany) with a 12-element Head Matrix coil, as described in an earlier publication^36^. The anatomical data were acquired using a Dual-Echo Turbo Spin Echo (PD-T2-TSE) sequence (TR 12070 ms; TE 13 ms and 102 ms) and a sagittal 3D-T1 Magnetization Prepared Rapid Acquisition Gradient Echo (MPRAGE) sequence (TR 1900 ms; TE 3.26 ms; inversion time 900 ms) with whole brain coverage. 1×1×1 mm^3^ isotropic resolution was used for both sequences.

Single shell diffusion-weighted data was acquired using a standard twice-refocused Spin Echo-Echo Planar Imaging (SE-EPI) sequence (FOV 208 mm; 64 slices; TR 9300 ms; TE 87 ms). 2×2×2 mm^3^ isotropic resolution was used for the sequence and the b-value was 1000 s/mm. There were in total 96 unique diffusion encoding directions in a three-part multi-scan DTI sequence. Each part consisted of uniformly distributed 31, 32 or 33 directions and three b0 images (images without diffusion encoding) that were taken in the beginning, in the middle, and in the end of each scan.

### 2.3. Imaging Analysis

#### 2.3.1. Striatal Segmentation

A study specific template was created from the structural MRI data. Then the regions of interest (ROIs) were manually labeled both on the template and the variants of the template to ensure accuracy. The data was clustered into 21 clusters using the Jacobian determinants from non-linear transformations derived from the template construction. Central-most subject of each cluster was chosen, and the study specific template was warped to them. The ROIs were manually segmented in the 21 versions of the same template. The segmentations were unwarped back to the standard template and final ROI labels were created. Detailed descriptions of the procedure have been provided in our prior articles ^37,38^.

#### 2.3.2. DTI Preprocessing

Good quality b0 images were chosen manually, coregistered, averaged, and moved in front of each 4D series. Brain masks were created based on the b0 volumes with the Brain Extraction Tool (BET) ^39^ by FSL (FMRIB Software Library v 5.0.9 ^40^. DTIPrep software ^41^ was used to inspect the quality of the data. Low quality diffusion images were discarded. Remaining images were also visually inspected, and more directions were excluded as needed. We have previously found that after the quality control steps, datasets that have more than 20 diffusion encoding directions will yield reliable tensor estimates ^42^. Here all infants with at least 20 diffusion encoding directions were selected and we used all available data thereafter. Eddy current and motion correction steps were conducted with FSL ^43^ and the b-vector matrix was rotated accordingly. A diffusion tensor model was fitted to each voxel included in the brain mask with the DTIFIT tool in FDT (FMRIB’s Diffusion Toolbox) of FSL. The DTI preprocessing steps have been provided in detail in our previous publication that also reports good test-retest repeatability in between segments of DTI sequences ^36^.

#### 2.3.3. Extracting Diffusion Metrics

To extract the MD values of the ROIs, the diffusion data were registered to the study-specific template space. This was carried out by rigidly registering the b0 images to nonuniformity-corrected T1-weighted data and combining the transformations.

from b0-to-T1 and the T1-to-template space for MD maps ^37,38^. The masking of the individual structures was then performed in the group average space to get the diffusion measures. We first defined the values from the anatomical masks in the template space. Second, to eliminate the partial volume effect, 1.5 mm erosion to the template masks ^44^. Mean MD values within each ROI was calculated with non-eroded masks and eroded masks. Here, the dataset created with values from the eroded, *i.e.* partial volume corrected, masks was used for the derived brain measures. The same analyses were repeated using the dataset created with values from the non-eroded masks and the results are presented in the supplementary materials (see Supplementary Material Figure S1 and Table S1 and S2).

### 2.4. Statistical Analysis

The associations between maternal pre-pregnancy BMI and the mean MD of the ROIs were investigated with a linear regression model that was adjusted for infant’s sex and postnatal age in days. The possible confounding effect of maternal GDM was considered ^45,46^. The same regression model was conducted in a subsample that included only the mothers without GDM. We performed sensitivity analyses that included other variables as covariates: infant’s birthweight and gestational age at birth in weeks, mother’s age at birth in years, socioeconomic status represented with education level (years of formal education) and the use of selective serotonin reuptake inhibitor, serotonin and norepinephrine reuptake inhibitor or other drugs that affect the central nervous system by individually adding these factors to and removing them from the regression model, to verify that the associations were not explained by these variables.

To assure the appropriateness of the regression models we checked the variance inflation factor (<1.5 for all variables in all models) and visually inspected the distribution of residuals and the Q-Q plots for normality.

Multiple comparisons were corrected with false discover rate (FDR) correction. *P* < 0.05 after FDR correction for multiple comparisons were considered statistically significant. All statistical analyses and data visualization were done in R 4.0.3 ^47^.

## 3. Results

### 3.1. Demographic Overview

Demographics of infants and mothers are presented in Table 1 and the mean MD values of each six regions or interest are presented in Table 2. Female infants had smaller mean MD values in the right putamen, however there were no statistically significant sex differences in mean MD values of infants in any ROI after correcting for multiple comparisons (Table 2).

**Table 1.** Demographics of infants and mothers presented with mean scores (range) or frequencies for the whole sample, male infants, and female infants separately. *P* values for the sex differences are shown in the far-right column.

| variable | whole sample<br>n = 116<sup>1</sup> | males<br>n = 66<sup>1</sup> | females<br>n = 50<sup>1</sup> | $p^2$ |
| --- | --- | --- | --- | --- |
| birth weight (g) | 3 482<br>(2 530 – 4 700) | 3 565<br>(2 720 – 4 700) | 3 390<br>(2 530 – 4 510) | 0.041 |
| birth height (cm) | 50.43<br>(44 – 56) | 50.74<br>(46 – 56) | 50.2<br>(44 – 53) | 0.048 |
| head circumference (cm) | 35.04<br>(32.5 – 37.5) | 35.27<br>(33 – 37.5) | 34.73<br>(32.5 – 37) | 0.04 |
| gestational weeks at birth | 39.88<br>(36.3 – 42.1) | 39.87<br>(37.6 – 42) | 39.9<br>(36.3 – 42.1) | 0.8 |
| gestational weeks at scan | 43.56<br>(41.1 – 46.4) | 43.55<br>(42 – 45.7) | 43.57<br>(41.1 – 46.4) | >0.9 |
| infant age at scan from birth (days) | 25<br>(8 – 45) | 25<br>(14 – 40) | 25<br>(8 – 45) | >0.9 |
| maternal pre-pregnancy body mass index | 24.2<br>(17.5 – 38.4) | 24.6<br>(18 – 38.4) | 23.8<br>(17.5 – 34.9) | 0.5 |
| underweight<br>BMI < 18 | 3 / 116 | 1 / 66 | 2 / 50 |  |
| normal weight<br>BMI 18-25 | 76 / 116 | 44 / 66 | 32 / 50 |  |
| overweight<br>BMI 25-30 | 25 / 116 | 12 / 66 | 13 / 50 |  |
| obese<br>BMI > 30 | 12 / 116 | 9 / 66 | 3 / 50 |  |
| gestational diabetes<br>mellitus |  |  |  | >0.9 |
| absent | 100 / 116 | 57 / 66 | 43 / 50 |  |
| present<br>(insulin<br>treatment) | 16 (1) / 116 | 9 (0) / 66 | 7 (1) / 50 |  |
| SSRI/SNRI<br>or other CNS<br>affecting drugs | 8 / 116 | 7 / 66 | 1 / 50 | 0.13 |
| maternal age at<br>birth (years) | 29.7<br>(20 – 41) | 30.1<br>(20 – 41) | 29.3<br>(21 – 41) | 0.3 |
| maternal education<br>level at birth |  |  |  | 0.8 |
| <12 years | 32 / 116 | 18 / 66 | 14 / 50 |  |
| 12 -15 years | 36 / 116 | 19 / 66 | 17 / 50 |  |
| 15 + years | 48 / 116 | 29 / 66 | 19 / 50 |  |

<sup>1</sup> Mean (range) or Frequency
<sup>2</sup> Wilcoxon rank sum test; Pearson's Chi-squared test

**Table 2.** Mean MD (SD) × 10^-3^mm^2^/s values of the six regions of interest (ROIs) for the whole sample, male and female infants separately. *P* values for the sex differences are shown in the far-right column.

| ROI | whole sample<br>n = 116<sup>1</sup> | males<br>n = 66<sup>1</sup> | females<br>n = 50<sup>1</sup> | *p*<sup>2</sup> |
| --- | --- | --- | --- | --- |
| right<br>putamen | 1.033<br>(0.030) | 1.038<br>(0.029) | 1.027<br>(0.030) | 0.045 |
| left<br>putamen | 1.033<br>(0.030) | 1.038<br>(0.029) | 1.029<br>(0.030) | 0.11 |
| right<br>globus<br>pallidus | 1.016<br>(0.030) | 1.018<br>(0.028) | 1.013<br>(0.032) | 0.4 |
| left<br>globus<br>pallidus | 1.011<br>(0.028) | 1.011<br>(0.029) | 1.011<br>(0.026) | >0.9 |
| right<br>caudate | 1.101<br>(0.037) | 1.101<br>(0.037) | 1.101<br>(0.039) | 0.7 |
| left<br>caudate | 1.112<br>(0.044) | 1.113<br>(0.044) | 1.112<br>(0.043) | 0.06 |

<sup>1</sup> Mean (standard deviation)
<sup>2</sup> Wilcoxon rank sum test

Infant age had strong negative associations with mean MD in all ROIs after FDR correction: the right putamen (*p* for infant’s age= 2.14e-10, partial R^2^ = 0.3), left putamen (*p* for infant’s age = 5.59e-11, partial R^2^ = 0.32), right globus pallidus (*p* for infant’s age = 0.000248, partial R^2^ = 0.11), left globus pallidus (p for infant’s age = 0.000529, partial R^2^ = 0.1), right caudate (*p* for infant’s age = 0.000524, partial R^2^ = 0.1) and left caudate (*p* for infant’s age = 3.5e-05, partial R^2^ = 0.14).

### 3.2. Associations between Maternal pre-pregnancy BMI and the Mean MD of Striatum

We found a positive association between maternal pre-pregnancy BMI and the mean MD in the left caudate nucleus (see Figure 1 and Table 3), adjusted for infant’s sex and age in days. Maternal pre-pregnancy BMI had no statistically significant association between the mean MD in the lentiform nuclei and the right caudate nucleus. The results remained unchanged in the sensitivity analyses.

**Figure 1.**
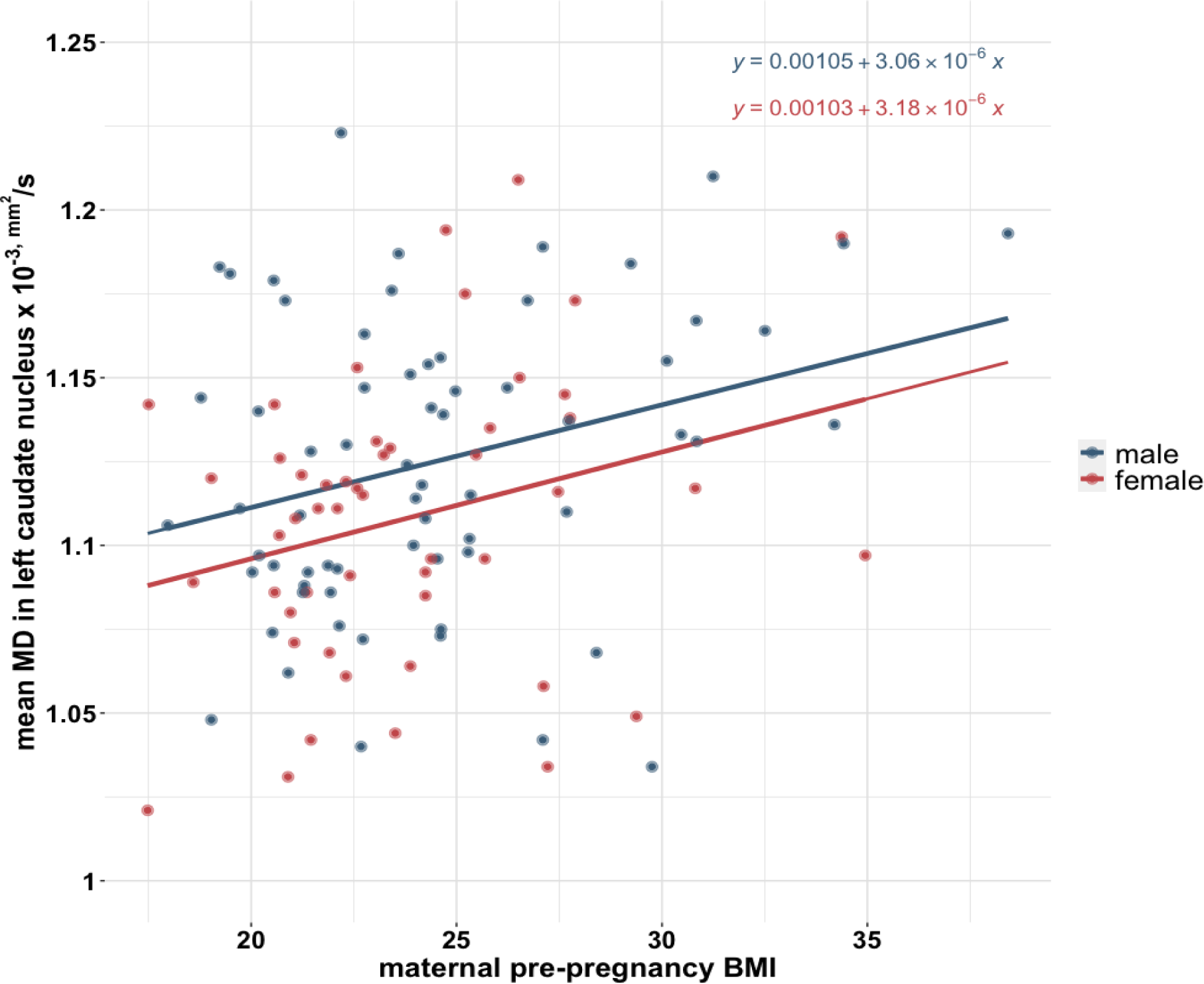
A scatterplot showing the association between maternal pre-pregnancy BMI and the mean mean diffusivity (MD) in left caudate nucleus.

**Table 3.**
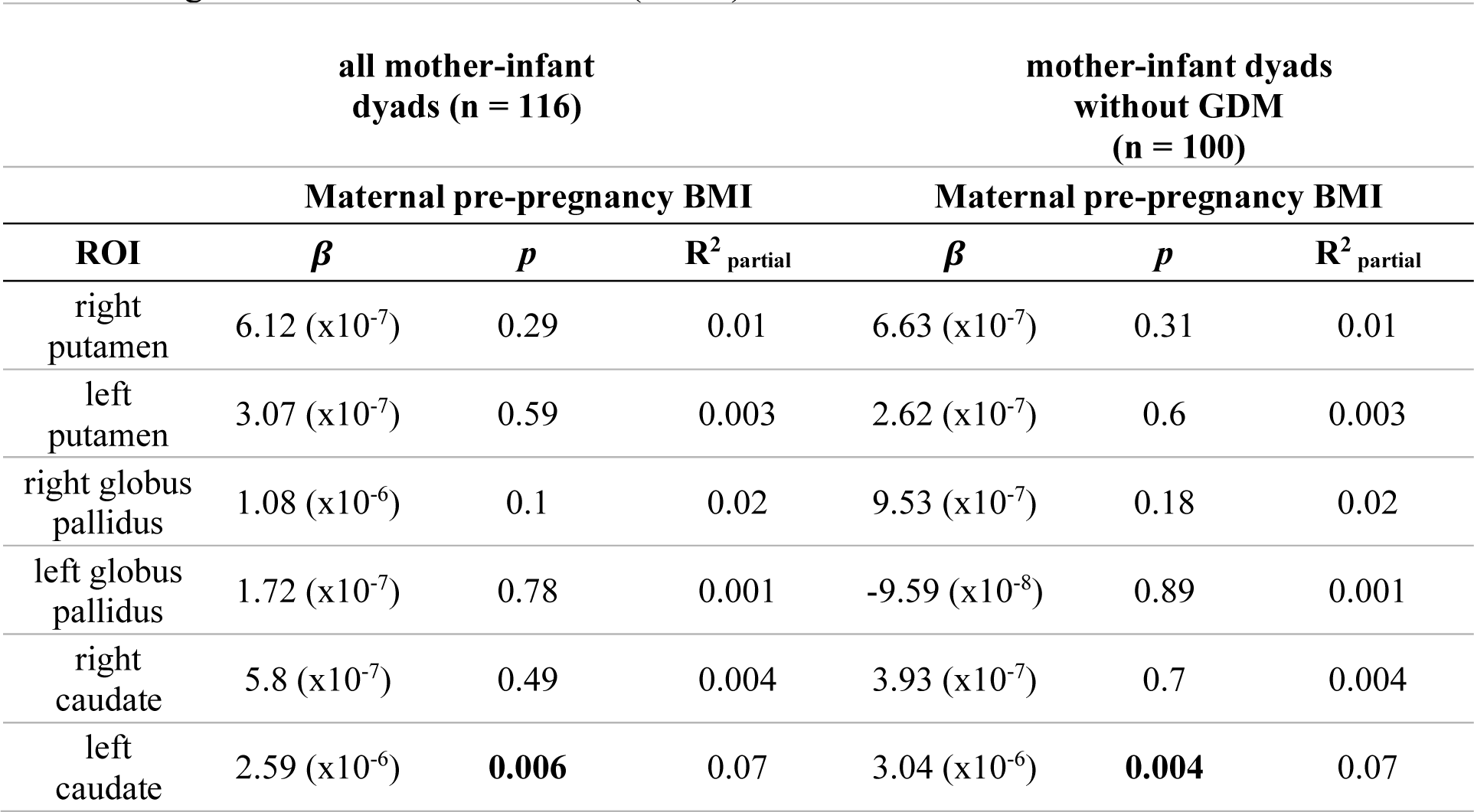
The association between maternal pre-pregnancy BMI and mean mean diffusivity (MD) of striatum adjusted for infant’s sex and age for the whole sample and the subsample without the gestational diabetes mellitus (GDM).

|  | all mother-infant<br>dyads (n = 116) |  |  | mother-infant dyads<br>without GDM<br>(n = 100) |  |  |
| --- | --- | --- | --- | --- | --- | --- |
|  | Maternal pre-pregnancy BMI |  |  | Maternal pre-pregnancy BMI |  |  |
| ROI | $\beta$ | *p* | $R^2_{\text{partial}}$ | $\beta$ | *p* | $R^2_{\text{partial}}$ |
| right putamen | $6.12 (x10^{-7})$ | 0.29 | 0.01 | $6.63 (x10^{-7})$ | 0.31 | 0.01 |
| left putamen | $3.07 (x10^{-7})$ | 0.59 | 0.003 | $2.62 (x10^{-7})$ | 0.6 | 0.003 |
| right globus pallidus | $1.08 (x10^{-6})$ | 0.1 | 0.02 | $9.53 (x10^{-7})$ | 0.18 | 0.02 |
| left globus pallidus | $1.72 (x10^{-7})$ | 0.78 | 0.001 | $-9.59 (x10^{-8})$ | 0.89 | 0.001 |
| right caudate | $5.8 (x10^{-7})$ | 0.49 | 0.004 | $3.93 (x10^{-7})$ | 0.7 | 0.004 |
| left caudate | $2.59 (x10^{-6})$ | **0.006** | 0.07 | $3.04 (x10^{-6})$ | **0.004** | 0.07 |

To assess the associations without the possible confounding effect of GDM, the cases with GDM were eliminated from the dataset (remaining n = 100) and the same linear regression model was applied to the subset. The results remained the same; maternal pre-pregnancy BMI was still significantly correlated with mean MD in the left caudate nucleus but not with other striatal brain regions.

## Discussion

This study applied an ROI analysis of DTI images to investigate the association between maternal pre-pregnancy BMI and newborn offspring striatal microstructural maturation. We found that maternal pre-pregnancy BMI was positively associated with left caudate MD. Moreover, this study provides conceptual advance for the investigation of *in utero* exposure to maternal adiposity before environmental factors such as diet, lifestyle or socio-economic statues start to contribute.

Reward sensitivity is a critical contributor to weight gain ^48^ as a behavioral modifier in promoting excess caloric intake. Increased reward sensitivity to food stimuli has been linked to excessive weight gain^18^. Indeed, increased caudate activation to high-sugar content food was repeatedly observed in high-risk adolescents due to parental obesity compared to low-risk adolescents^49,50^. It can thus be speculated that the involvement of the caudate nucleus might be partly defined from the *in utero* exposure to maternal obesity. By showing an association between maternal BMI and infant left caudate MD, the current study gives support to this hypothesis. Future longitudinal studies are critical to assess whether there is a programming effect.

The association between maternal pre-pregnancy BMI and neonate striatal MD was asymmetrical. Although it is important to note that the lack of bilateral findings may simply due to the high amount of noise present in newborn MRI ^51^. However, in a deep learning study where the researchers used the anatomical scans of almost 18,000 adult participants to create a convolutional neural network to predict the BMI with high accuracy, it was found that left caudate had a strong influence on predicting the BMI ^52^. This could imply that true hemispheric asymmetry is indeed present in the studied mechanisms. Moreover, an fMRI study showed that adolescents at high-risk for obesity (due to paternal obesity) showed larger right caudate responses to sugary food cues compared to the low-risk group ^54^. Structural asymmetry and functional lateralization of the potential impact of parental BMI should be studied more to draw any further conclusions. The same applies to the possible differences between *in utero* exposure to maternal obesity and exposure later in life, to gain a better understanding of the neural mechanisms involved.

The microstructural integrity of the putamen, more than any other ROI in the study, was explained largely by the age of the infants in days. Previous research revealed that the putamen is one of the fastest growing regions in the infant brain, and is assumed to have high metabolic activity ^55^. Another study showed that vessel density in the putamen rapidly increases after birth ^56^. The strong association we found with infant’s age and putamen MD is in line with previous studies and provides a more complete understanding of the neurodevelopment of infant gray matter.

As opposed to the previous findings showing lowered MD values in putamen and globus pallidus in mildly obese adults^28^, no such association was found with maternal adiposity and striatal MD values in neonates. However, as speculated in the related paper the association could be explained by the number of the dopamine receptors in the striatal area. Takeuchi et al. argued that excessive hedonic eating, is related to the increased number of dopamine receptors in the striatal areas and the striatal axonal density is observed as lowered MD (2020). It is important to highlight the importance of future DTI and positron emission tomography multimodal imaging studies to investigate the underlying neural mechanisms of *in utero* exposure to obesity.

Here we present on an adverse effect of maternal pre-pregnancy BMI on the striatal development that may represent a key mechanism of offspring weight gain from birth to adulthood. Because it is well-established that primary prevention of obesity is more effective than treating adult obesity^57,58^, the investigation of alterations driven by maternal pre-pregnancy BMI in the offspring’s reward system is of high relevance to improving offspring obesity-related outcomes by planning effective interventive measures ^59^.

Furthermore, research shows that the adverse effect of maternal obesity on neurodevelopment manifests on a broader spectrum than problems in energy balance, including various cognitive impairments and psychiatric disorders ^60,61^. Specifically, a large meta-analysis conducted with the data of 36 cohorts from several European countries and the United States of America revealed that maternal obesity is posing an increased risk for attention and hyperactivity disorder, autism spectrum disorders, and cognitive delay^62^. However, the causality is still speculative^63^ and may be informed through the identification of mechanistic pathways such as fetal programming of reward circuitry. Thus, more research is warranted to further consider the extent to which adverse effects of *in utero* exposure to maternal BMI have on the physiological and mental well-being of future generations.

A limitation of the current study is small number of participants with GDM. A larger sample of mothers with GDM would have provided the possibility of investigating and perhaps comparing the impact of GDM alongside with BMI. Furthermore, our whole sample was drawn from Finnish population. Similar studies with participants from more ethnically diverse backgrounds are needed to improve generalizability of the findings.

## Conclusion

The findings of this study suggest a positive association between maternal pre-pregnancy weight status and the mean MD values of the left caudate nucleus. As increased MD is commonly interpreted as lower tissue density, it could be concluded that *in utero* exposure to maternal obesity might have a growth / maturation impairing impact on the tissue density of infant left caudate nucleus. Further investigations are required to identify the prognostic relevance of early life caudate development and weight trajectories of the offspring who were born to obese mothers.

## Declarations

### Ethics Approval and Consent to Participate

This study was conducted in accordance with the Declaration of Helsinki, and it was approved by the Ethics Committee of the Hospital District of Southwest Finland (15.03.2011) §95, ETMK: 31/180/2011.

### Consent for Publication

Not applicable.

### Availability of Data and Materials

The current Finnish legislation and our Ethical Board approval do not permit the open data sharing of the imaging data or derived measures. Investigators interested in getting access to the data are encouraged to contact FinnBrain’s Principal Investigators (https://sites.utu.fi/finnbrain/en/contact/). The analysis code can be made available upon a reasonable request to the corresponding author.

### Competing Interests

The authors declare no competing interests.

## Acknowledgements

Krisse Kuvaja and Satu Lehtola for scanning the infants. Maria Keskinen for recruiting the participants.

## List of Abbreviations

SD: standard deviation
BMI: body mass index
DTI: diffusion tensor imaging
MD: mean diffusivity
MRI: magnetic resonance imaging
GDM: gestational diabetes mellitus
PD-T2-TSE: dual-echo turbo spin echo
TR: repetition time
TE: echo time
MPRAGE: magnetization prepared rapid acquisition gradient echo
SE-EPI: spin echo - echo planar imaging
FOV: field of view
ROI: region of interest
BET: brain extraction tool
FSL: FMRIB software library
FDR: false discovery rate

---

**We report here the regression results that were obtained without performing a partial volume correction to the anatomical labels as stated in the main manuscript:** “The masking of the individual structures was then performed in the group average space to get the diffusion measures. We first defined the values from the anatomical masks in the template space. Second, to eliminate the partial volume effect, 1.5 mm erosion to the template masks ^42^. Mean MD values within each ROI was calculated with non-eroded masks and eroded masks. Here, the dataset created with values from the eroded, *i.e.* partial volume corrected, masks was used for the derived brain measures. The same analyses were repeated using the dataset created with values from the non-eroded masks and the results are presented in the supplementary materials (see Supplementary Material Figure S1 and Table S1 and S2).”

**The overall pattern of results is identical although differences are also apparent.**

**Figure S1.**
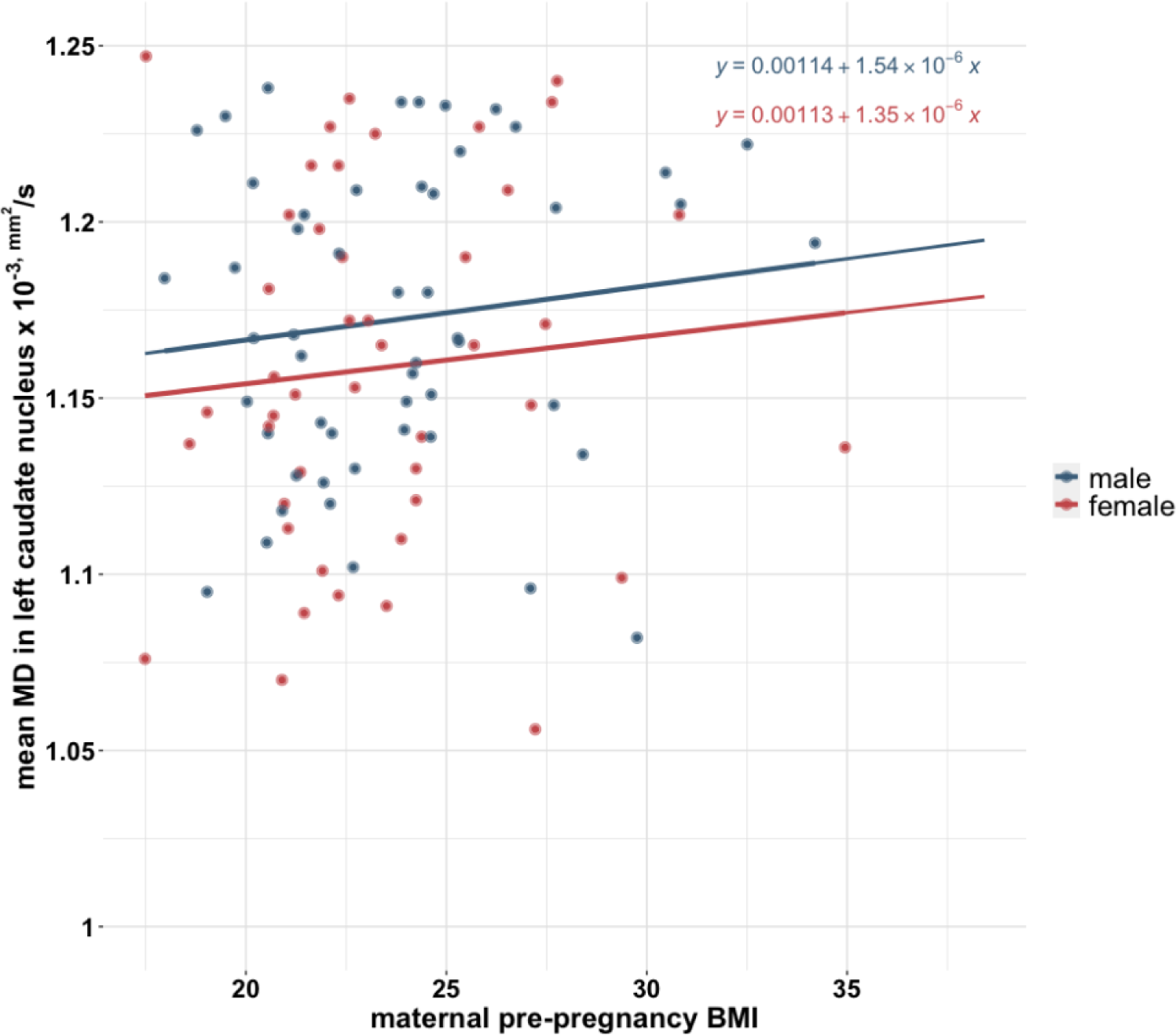
A scatterplot showing the association between maternal pre-pregnancy BMI and the mean MD in left caudate nucleus.

**Table S1.**
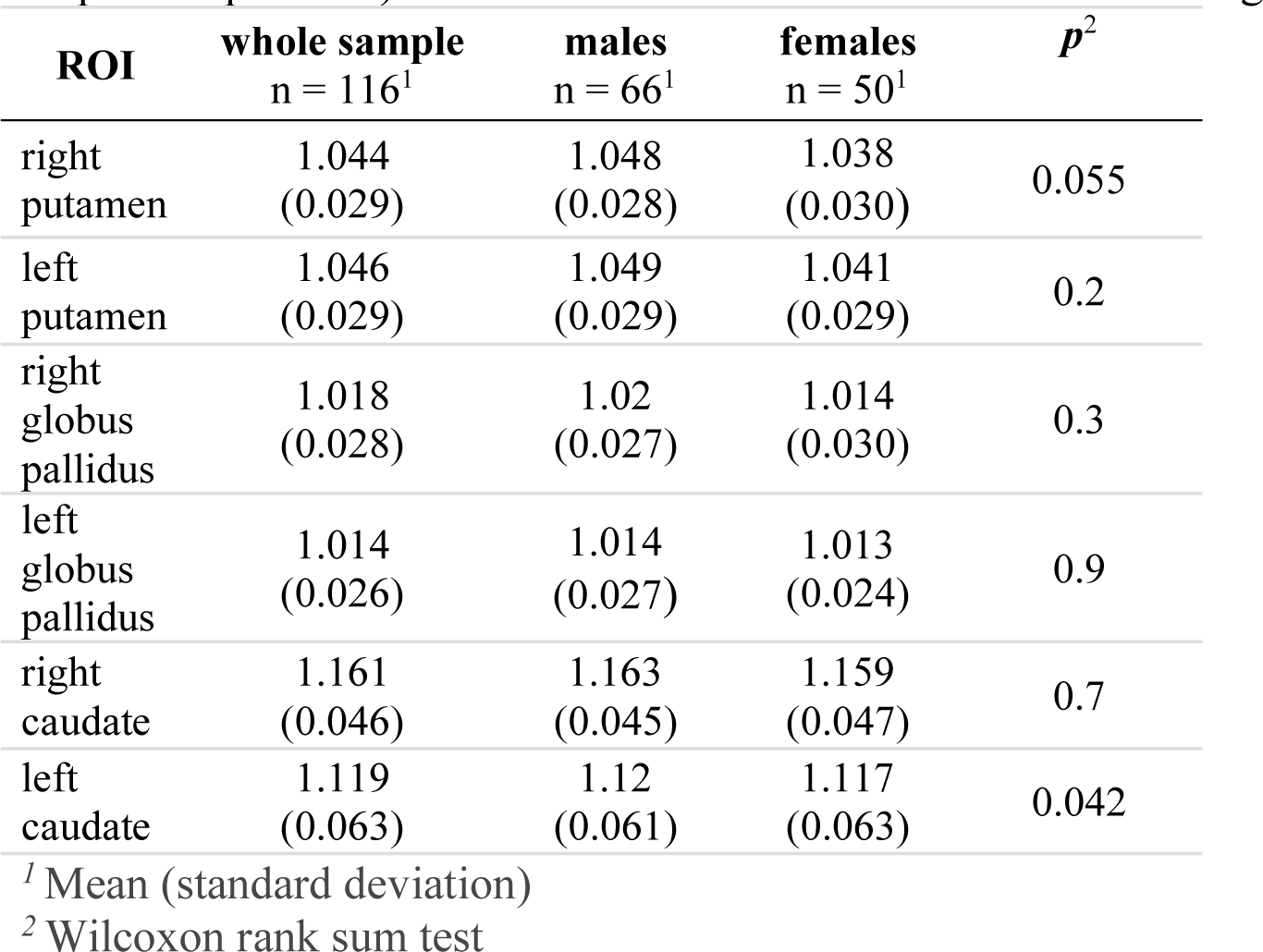
Mean MD (SD) × 10^-3^mm^2^/s values of the six regions of interest (ROIs) for the whole sample, male and female infants separately, using the non-eroded data. *P* values (not corrected for multiple comparisons) for the sex differences are shown in the far-right column. A significant positive association was found between maternal pre-pregnancy BMI and the mean MD in the left caudate nucleus (see Figure S1 and Table S2). The results remained unchanged in the sensitivity analyses.

| ROI | whole sample<br>n = 116<sup>1</sup> | males<br>n = 66<sup>1</sup> | females<br>n = 50<sup>1</sup> | *p*<sup>2</sup> |
| --- | --- | --- | --- | --- |
| right putamen | 1.044<br>(0.029) | 1.048<br>(0.028) | 1.038<br>(0.030) | 0.055 |
| left putamen | 1.046<br>(0.029) | 1.049<br>(0.029) | 1.041<br>(0.029) | 0.2 |
| right globus pallidus | 1.018<br>(0.028) | 1.02<br>(0.027) | 1.014<br>(0.030) | 0.3 |
| left globus pallidus | 1.014<br>(0.026) | 1.014<br>(0.027) | 1.013<br>(0.024) | 0.9 |
| right caudate | 1.161<br>(0.046) | 1.163<br>(0.045) | 1.159<br>(0.047) | 0.7 |
| left caudate | 1.119<br>(0.063) | 1.12<br>(0.061) | 1.117<br>(0.063) | 0.042 |

<sup>1</sup> Mean (standard deviation)
<sup>2</sup> Wilcoxon rank sum test

**Table S2.**
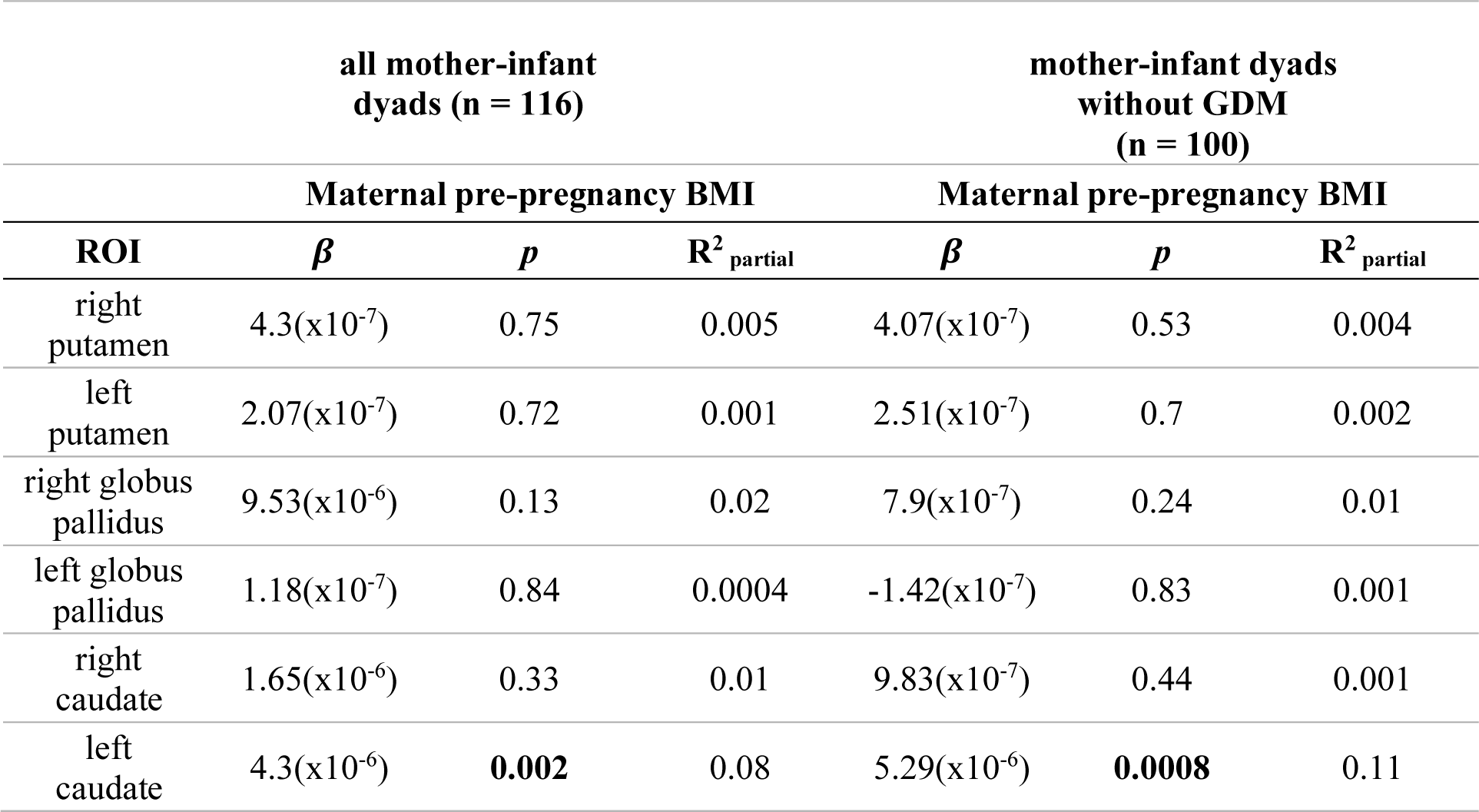
The association between maternal pre-pregnancy BMI and mean mean diffusivity of striatum adjusted for infant’s sex and age for the whole sample and the subsample without the gestational diabetes mellitus (GDM).

|  | all mother-infant dyads (n = 116) |  |  | mother-infant dyads without GDM (n = 100) |  |  |
| --- | --- | --- | --- | --- | --- | --- |
|  | Maternal pre-pregnancy BMI |  |  | Maternal pre-pregnancy BMI |  |  |
| ROI | $\beta$ | *p* | $R^2_{\text{partial}}$ | $\beta$ | *p* | $R^2_{\text{partial}}$ |
| right putamen | $4.3(\times 10^{-7})$ | 0.75 | 0.005 | $4.07(\times 10^{-7})$ | 0.53 | 0.004 |
| left putamen | $2.07(\times 10^{-7})$ | 0.72 | 0.001 | $2.51(\times 10^{-7})$ | 0.7 | 0.002 |
| right globus pallidus | $9.53(\times 10^{-6})$ | 0.13 | 0.02 | $7.9(\times 10^{-7})$ | 0.24 | 0.01 |
| left globus pallidus | $1.18(\times 10^{-7})$ | 0.84 | 0.0004 | $-1.42(\times 10^{-7})$ | 0.83 | 0.001 |
| right caudate | $1.65(\times 10^{-6})$ | 0.33 | 0.01 | $9.83(\times 10^{-7})$ | 0.44 | 0.001 |
| left caudate | $4.3(\times 10^{-6})$ | **0.002** | 0.08 | $5.29(\times 10^{-6})$ | **0.0008** | 0.11 |

## Notes

### Competing Interest Statement

The authors have declared no competing interest.

